# Daily dynamics of resting-state EEG theta and gamma fluctuations are associated with cognitive performance in healthy aging

**DOI:** 10.1101/2023.12.21.572745

**Authors:** Kenza Bennis, Francis Eustache, Fabienne Collette, Gilles Vandewalle, Thomas Hinault

**Affiliations:** Inserm, U1077, EPHE, UNICAEN, Normandie Université, PSL Université Paris, CHU de Caen, GIP Cyceron, Neuropsychologie et Imagerie de la Mémoire Humaine (NIMH), 14000 Caen, France; GIGA-CRC-In Vivo Imaging, Université de Liège and Belgian National Fund for Scientific Research, Liège, Belgium

**Author notes:** Corresponding author: Thomas Hinault, INSERM-EPHE-UNICAEN U1077, 2 rue des Rochambelles, 14032 Caen, FRANCE. **Declarations of interest**: none.

**Keywords:** EEG fluctuations, daily dynamics, theta, gamma, cognitive functions, inter-networks

## Abstract

Healthy age-related cognitive changes are highly heterogeneous across individuals. This variability is increasingly explained through the lens of spontaneous fluctuations of brain activity, now considered as powerful index of age-related changes. However, brain activity is a biological process modulated by circadian rhythms, and how these fluctuations evolve throughout the day is under investigated. Assessing the daily dynamics of brain fluctuations involves the use of techniques measuring the temporal dynamics of synchronized communications between brain regions, such as electroencephalography. We found that theta and gamma daily fluctuations in the salience-control executive inter-network (SN-CEN) are associated with distinct mechanisms underlying cognitive heterogeneity in aging. Higher levels of SN-CEN theta daily fluctuations appear to be deleterious for memory performance and were associated with higher tau/neuroinflammation rates. In contrast, higher levels of gamma daily fluctuations are positively associated with executive performance, and were associated with lower rate of β-amyloid deposition. Thus, accounting for daily EEG fluctuations of brain activity contributes to better understand subtle brain changes underlying individuals’ cognitive performance in healthy aging.

## 1. Introduction

There is large interindividual variability in human cognitive performance, and performance also fluctuate across the lifespan at the individual level. Evidence exists suggesting that cognitive heterogeneity through human development follows a U-shaped trajectory, with young children showing the greatest heterogeneity, followed by least heterogeneity in young adulthood, and again larger heterogeneity when people grow old (Williams et al., 2005). During aging, some healthy individuals can show slight cognitive decline and are considered as potentially at risk of neurodegenerative pathologies, while others show similar cognitive performance to that of younger individuals (Hultsch et al., 2008). The brain bases of inter-individual variability in cognitive aging remain, however, only partly elucidated.

Fluctuation of brain activity follows an inverted U-shaped curve, such that, compared to young adults, young children and older adults show lower fluctuations in BOLD-signal (Garrett et al., 2011) and EEG signal (Sleiman-Malkoun et al., 2015). These spontaneous fluctuations of brain activity were first considered to consist of noise, but are now known to critically contribute to brain function for optimal brain responsivity to changes in the environment (Uddin et al., 2020; Garrett et al., 2014). Thus, recent imaging studies aim to better understand individual cognitive trajectories in aging through the lens of brain fluctuations (for review, see Waschke et al., 2021). Part of these works documented brain activity fluctuations in resting-state conditions, as it was assumed that brain activity at rest in so-called Resting-State Networks (Power et al., 2011, Yeo et al., 2011), reflect the stable and intrinsic functional connectivity of the brain that is linked with cognitive abilities (Miraglia et al., 2017; Nashiro et al., 2017, Engel et al., 2021). Reduced fluctuations of brain functional connectivity in elderly were associated to lower cognitive performance relative to young adults, while higher fluctuations of brain functional connectivity were interpreted as reflecting preserved brain communication efficiency and flexibility in aging (Kumral et al., 2020; Grady et Garrett, 2018). These findings provide arguments for further investigating brain fluctuations at rest to shed new lights on the cognitive heterogeneity in healthy aging.

Brain rhythms and their fluctuations are increasingly investigated through time-frequency methods, that decompose the signal in five frequency bands, namely from the slowest to the fastest rhythms: delta (1-4Hz), theta (4-8Hz), alpha (8-12Hz), beta (13-30Hz) and gamma (30-100Hz). EEG functional connectivity can be investigated through the phase synchrony of these brain rhythms between distant brain regions, which provides information about cognitive functioning in aging (see Babiloni et al., 2020 for a review). Relative to young adults, a global slowing of brain activity was observed with aging, reflected by an increase of theta and delta phase synchrony and a decrease of beta and gamma phase synchrony, which has been associated with lower cognitive performance (Lopez et al., 2014, Vecchio et al., 2014). Interestingly, older adults with cognitive performance similar to younger individuals exhibit an increase in brain synchrony between distant brain regions, interpreted as a compensatory mechanism (Ariza et al., 2015; Frutos Lucas et al., 2020). Noteworthy, normal aging is associated with the presence of tau and β-amyloid proteins, which abnormal accumulation can lead to pathological cognitive decline (Hanseeuw et al.,2019). Impairments in phase synchrony between distant regions underlying specific cognitive processes, occur long before age-related structural alterations or β-amyloid and tau proteins abnormal deposition (Babiloni et al., 2020, 2021). Therefore, EEG signal is a direct marker of brain activity that allows better detection of acute age-related brain changes and their interpretation in regard to individuals’ cognitive performance.

In addition, to vary with aging and across individuals, cognition is not stable across the day (Valdez, 2019; Waterhouse, 2010). Cognitive performance depends on the interplay between prior sleep-wake history, which sets the need for sleep, and the circadian system which promotes wakefulness and cognition during the day, and favors sleep at night (Gaggioni et al., 2019). This interplay also results in daily fluctuations in brain response characterized as cortical excitability, that were linked to cognitive changes in aging, with large fluctuation associated with preserved cognitive abilities in healthy older adults (Van Egroo et al. 2019, Gaggioni et al. 2019). However, resting-state connectivity assessed by EEG time-frequency studies have mostly investigated phase synchrony of brain rhythms fluctuations through short single recordings of approximately 5 minutes. Thus, how fluctuation in phase synchrony of brain rhythms evolve throughout the day at resting in healthy aging and how they are associated with cognitive performance remain insufficiently characterized. We reasoned that the fluctuation of phase synchrony should be related to cognitive performance as well as tau and β-amyloids burdens, and could provide novel markers of age-related cognitive decline.

We therefore investigated the variability of brain activity synchrony within resting-state networks across the day in healthy late middle-aged participants aged 50 to 70 years old. Our goals were threefold: i) investigate brain activity synchrony fluctuations across the day at rest in healthy aging: we expected more synchrony fluctuations among networks at the beginning compared to the end of the day, thus reflecting the daily dynamics of brain rhythms fluctuations; ii) study the relationships between global daily brain rhythms synchrony fluctuations and cognitive performance: we hypothesized that high level of fluctuations during the day would be associated with better cognitive performance, thus reflecting preserved brain efficiency to maintain cognitive abilities; iii) assess the association between global brain rhythms synchrony fluctuations across the day and pathological markers: we expected that high amount of tau/neuroinflammation would be associated with lower global daily brain synchrony fluctuations, as it would reflect a deleterious influence on brain functioning and communication.

## 2. Methods

### 2.1. Experimental protocol

Participants were enrolled in a multimodal longitudinal study designed to identify cerebral biomarkers of normal cognitive aging (the Cognitive Fitness in Aging – COFITAGE – study; Van Egroo et al., 2019). Five EEG recordings of spontaneous resting-state activity were performed on the same day, between 10a.m. and 1a.m. in the context of a 20-hour continuous wakefulness protocol under strictly controlled constant routine conditions (i.e., in-bed semi-recumbent position, dim light <5 lux, temperature ∼19°C, regular isocaloric food intake, no time-of-day information and sound-proofed rooms). Neuropsychological assessment, β-amyloid-PET and Tau/neuroinflammation-PET imaging together with T1-weighted MRI were also acquired on separate visits. All procedures were previously reported (first in Van Egroo et al., 2019, 2021; Narbutas et al., 2019,2021; Chylinski et al., 2021, 2022; Rizzolo et al., 2021) (Figure 1A).

**Figure 1.**
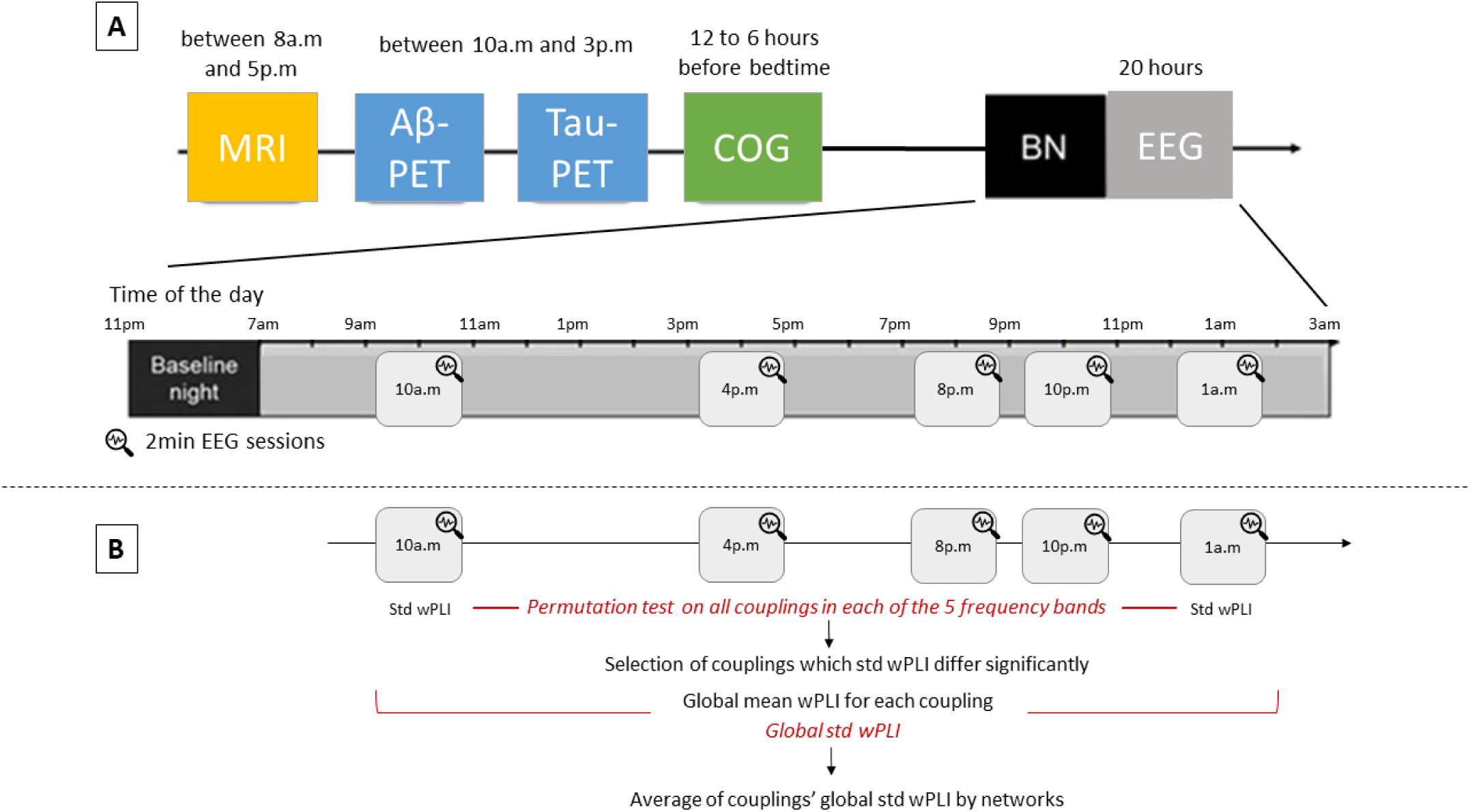
**A**. Experimental protocol, adapted from Van Egroo et al. (2019). COG: cognitive assessment, BN: Baseline-Night. **B.** Schematic representation of statistical analyses on EEG signal. Std wPLI: standard deviation of the weighted phase lag index (wPLI).

### 2.2. Participants

We analyzed data from one hundred and one healthy late middle-aged participants (68 women and 33 men; aged 50-69 years). No participants reported any recent history of neurological or psychiatric disease or were taking medication affecting the central nervous system. Extended information about protocol, exclusion criteria, recruitment, consent and financial reward can be found in previous publications (Van Egroo et al., 2019; Chylinski et al., 2021, 2022; Narbutas et al., 2021). A subsample of 64 participants who had data for tau/neuroinflammation-PET imaging was also considered for additional analyses. Demographic characteristics of the final samples are described in Table 1. The study was approved by the Ethics Committee of Medicine Faculty of Medicine of the University of Liège, Belgium.

**Table 1.** Demographics, cognitive scores, tau and BA burden, structural and functional measures for the entire sample.

| NEUROPSYCHOLOGICAL ASSESSMENT |  |
| --- | --- |
| NUMBER OF PARTICIPANTS | 101 |
| NUMBER OF WOMEN, N (%) | 68 (67.3%) |
| AGE | 59.4 (5.287) |
| YEARS OF EDUCATION | 15.2 (3.009) |
| MEMORY COMPOSITE SCORE | 0.014 (0.925) |
| MEMORY SUBTEST: MST (RECOGNITION MEMORY SCORE) | 0.792 (0.153) |
| EXECUTIVE COMPOSITE SCORE | -0.038 (0.908) |
| ATTENTION COMPOSITE SCORE | 0.039 (0.981) |
| AB AND TAU/NEUROINFLAMMATION – PET IMAGING |  |
| TAU/NEUROINFLAMMATION BURDEN |  |
| NUMBER OF PARTICIPANTS | 64 |
| NUMBER OF FEMALES, N (%) | 42 (65.6%) |
| THK-PET IN BRAAK I/II ROIS (SUVR) | 2.289 (0.247) |
| B-AMYLOID BURDEN |  |
| NUMBER OF PARTICIPANTS | 101 |
| NUMBER OF FEMALES, N (%) | 68 (67.3%) |
| B-AMYLOID EARLY STAGE (CL) | 0.855 (0.052) |
| MRI |  |
| CORTICAL GRAY MATTER VOLUME (MM<sup>3</sup>) | 458647.230 (39834.230) |
| PHASE SYNCHRONY |  |
| GLOBAL STD WPLI THETA – SN-CEN | 0.046 (0.021) |
| GLOBAL STD WPLI GAMMA – SN-CEN | 0.029 (0.013) |

### 2.3. Neuropsychological assessment

Neuropsychological assessment was administered in two 1.5h sessions and consisted of a battery of cognitive tests assessing three specific domains: memory, attention and executive functions (Table 1; see supplementary for the entire set of neuropsychological assessment). The raw scores were converted to z-scores and three domain-specific composite scores were computed as the standardized sum of z-scores of the domain-specific scores, where higher values indicate better performance. We focused our analyses on the four composite scores and we also included the recognition memory score of the Mnemonic Similarity task (MST), as a previous published work on the COFITAGE database (Rizzolo et al., 2021), in line with the literature (Pishdadian et al., 2020) showed that it might be an early cognitive marker of memory decline among the group. A detailed description of the neuropsychological assessment can be found in previous work (Narbutas et al., 2019; Van Egroo et al., 2019).

### 2.4. PET imaging

β-Amyloid-PET and Tau/neuroinflammation-PET imaging were performed on an ECAT EXACT+ HR scanner (Siemens, Erlangen, Germany). β-Amyloid-PET imaging was performed with radiotracers [18F]Flutemetamol except for three subjects for which [18F]Florbetapir was used. Tau/neuroinflammation-PET imaging was performed with [18F]THK5351 for all subjects. For both β-Amyloid and tau/neuroinflammation PET imaging, a standardized uptake value ratio (SUVR) was calculated (Table 1). As β-Amyloid-PET imaging were acquired using different radioligands, their SUVR values were converted into Centiloid units (Klunk et al., 2015) (in line with previous works of this cohort, see Narbutas et al., 2019 and Van Egroo et al., 2019). Volumes of interest were determined using the automated anatomical labeling atlas (AAL) (Tzourio-Mazoyer et al., 2002). β-Amyloid burden was averaged over composite masks covering neocortical regions reported to undergo the earliest aggregation sites for β-Amyloid pathology (Grothe et al., 2017), while Tau/neuroinflammation burden was averaged over regions corresponding to Braak stages of early regional tau pathology (Braak and Braak, 1991; Shöll et al., 2016). The detailed PET imaging procedure was previously published in Narbutas et al., 2021.

### 2.5. Anatomical data

Participants’ T1-weighted MRI acquisition was performed on a 3-Tesla MR scanner (MAGNETOM Prisma, Siemens) to assess brain grey matter integrity (Table 1). The following parameters were used: repetition time (TR) = 18.7ms; flip angle (FA) = 20 degrees; 3D multiecho fast low angle shot (FLASH) sequence (TR/FA) = 136166; voxel size = 1 mm^3^ isotropic; acquisition time = 19minutes (see Van Egroo and al., 2019 for detailed parameters). The FreeSurfer (Fischl, 2012) software was used to generate cortical surfaces and automatically segment cortical structures from each participant’s T1-weighted anatomical MRI, to account for individual brain anatomy during source reconstruction.

### 2.6. EEG recording and analyses

#### 2.6.1. Data acquisition

For each participant, two minutes of resting-state EEG (sampling rate: 1450 Hz, bandpass filter: 0.1-500 Hz) were recorded five times throughout the wake-extension protocol (at 10:00 a.m., 4:00 p.m., 8:00 p.m., 10:00 p.m. and 1:00 a.m.) with a 60-channel EEG system (Eximia, Nexstim, Helsinki, Finland) covering the whole scalp. Participants were instructed to relax and avoid blinking while staring at a black dot.

#### 2.6.2. Pre-processing

Artifact and channel rejection (on continuous data), filtering (0.5-40Hz bandpass, on unepoched data), re-referencing (i.e., using the algebraic average of the left TP9 and right TP10 mastoid electrodes) and source estimation were performed using Brainstorm (Tadel et al., 2011). Physiological artefacts (blinks, saccades) were identified and manually removed through Independent Component Analyses (ICA; Comon, 1994) using Infomax algorithm (EEGLAB, runica.m). Independent Component Analyses approach consists in removing artifacts from the recording without removing the affected data portions, by identifying spatial components that are independent in time and uncorrelated with each other (Tadel et al., 2011).

#### 2.6.3. Sources reconstruction

FreeSurfer (Fischl, 2012) segmentation of individuals T1-weighted anatomical MRI was used to improve the accuracy of the source reconstruction, and to account for anatomical changes with age. The EEG forward model was obtained from a symmetric boundary element method (BEM model; OpenMEEG, Gramfort et al., 2010), fitted to the spatial positions of each electrode. A cortically constrained sLORETA procedure (Pascual-Marqui and Lehmann, 1994) was applied to estimate the cortical origin of scalp EEG signals. The estimated sources were then projected into a standard space (i.e., ICBM152 template) for comparisons between groups and individuals, while accounting for differences in native anatomy.

#### 2.6.4. Analyses

The individual alpha-peak frequency (IAF) observed at occipital sites was used to estimate the range of each frequency band. Based on previous works (Toppi et al., 2018) the following frequency bands were considered: Delta (IAF-8/IAF-6), Theta (IAF-6/IAF-2), Alpha (IAF-2/IAF+2), Beta (IAF+2/IAF+14) and Gamma1 (IAF+15/IAF+30). Phase-lag index (weighted PLI analyses; Stam et al., 2007, Vinck et al., 2011) was used to assess the functional synchrony between regions of interest (ROI) defined by using the Desikan-Kiliany atlas brain parcellation (Desikan et al., 2006). PLI analyses estimate the variability of phase differences between two regions over time. Similar phase difference across time is indicated by a PLI value close to 1 (i.e., high synchrony between regions), while large variability in the phase difference is indicated by a PLI value close to zero. PLI measure has been shown to be less sensitive to the influence of common sources and amplitude effects relative to phase-locking value, as it disregards zero phase lag that could reflect volume conduction artefacts (Stam et al., 2007, Hardmeier et al., 2014).

### 2.7. Statistical tests

Analyses were first conducted on the fluctuation rate (standard deviation across a resting-state session) of the wPLI calculated within each coupling (68×68 ROIs matrix, 4624 couplings) in each frequency band. Data were analyzed through permutation t-tests (with false discovery rate correction for multiple comparisons, FDR) performed in Brainstorm (Tadel et al., 2011), using a method originally implemented in Fieldtrip (Oostenveld et al., 2011). In each frequency band, we selected couplings for which the fluctuation rate of the wPLI differed significantly between the first and the fifth recordings.

For each of those couplings, we calculated global wPLI fluctuation rates across the five sessions (standard deviation of the global mean wPLI of the five sessions). Couplings’ global wPLI fluctuation rates were averaged within same intra-or inter-networks, according to their ROIs’ membership of the six Resting State Networks defined by Yeo et al. (2011) and labelled using their anatomical terminology in a draft network taxonomy by Uddin et al. (2019): Central Executive Network (CEN) i.e. Lateral Frontoparietal Network (L-FPN), Salience Network (SN) i.e. Midcingulo-Insular Network (M-CIN), Default Mode Network (DMN) i.e. Medial Frontoparietal Network (M-FPN), Dorsal Attention Network (DAN) i.e. Dorsal Frontoparietal Network (D-FPN), Visual System (VS) i.e. Occipital Network (ON) and Sensorimotor Network (SMN) i.e. Pericentral Network (PN) (Figure 1B) (see supplementary for detailed Desikan-Killany ROIs’ attribution to the six Resting State Networks).

Finally, regressions analyses were conducted to assess the association between networks’ global wPLI fluctuation rates, cognitive performance scores (the three domain-specific composite scores and the recognition memory score of the Mnemonic Similarity Task, MST), and β-amyloid and tau/neuroinflammation rates, using JASP (https://jasp-stats.org/; version 0.18.1). Participants’ age, sex and mean gray matter volume were included as covariates in the analyses. Results were FDR corrected for multiple comparisons (Benjamini and Hochberg, 1995) (see supplementary for detailed results concerning FDR correction).

## 3. Results

### 3.1. Theta and gamma inter-network connectivity fluctuates across the day

We first considered the changes in connectivity between the first and last session of the protocol (Figure 1B). Permutation t-test for each frequency band showed a significant increase of wPLI fluctuation rates between the first and the fifth recordings for inter-network couplings of the SN-CEN (t = –2.7999, p = 0.005) and of the DMN-CEN (t = –2.5178, p = 0.005) in the theta band, and a significant decrease for wPLI fluctuation rates for inter-network couplings of the SN-CEN (t = 2.8671, p = 0.005), of the DMN-SN (t = 2.5653, p = 0.005), of the CEN-ON (t = 2.8974, p = 0.005) and of the CEN-DAN (t = 2.6007, p = 0.005) in the gamma band. Across the day, theta fluctuations increase while they decrease in the gamma bands, especially in inter-network couplings. For both theta and gamma, we observed qualitatively a peak of fluctuation rates at the third recording (8p.m.).

As an index of inter-networks’ global synchrony fluctuation rates in a day, we averaged the global wPLI fluctuation rates across the five sessions, computed for each of their respective couplings (Figure 1B). To test the hypothesis that daily neural fluctuations could help explaining the heterogeneity of cognitive performance across older individuals, we then assessed the associations between global wPLI fluctuation rates of these identified inter-networks and the cognitive performance measured by the three-domain-specific composite scores and the recognition memory score. Results were therefore focused on SN-CEN inter-network which was the only one associated to cognitive performance in both theta and gamma bands in the sections to follow (Figure 2A).

**Figure 2.**
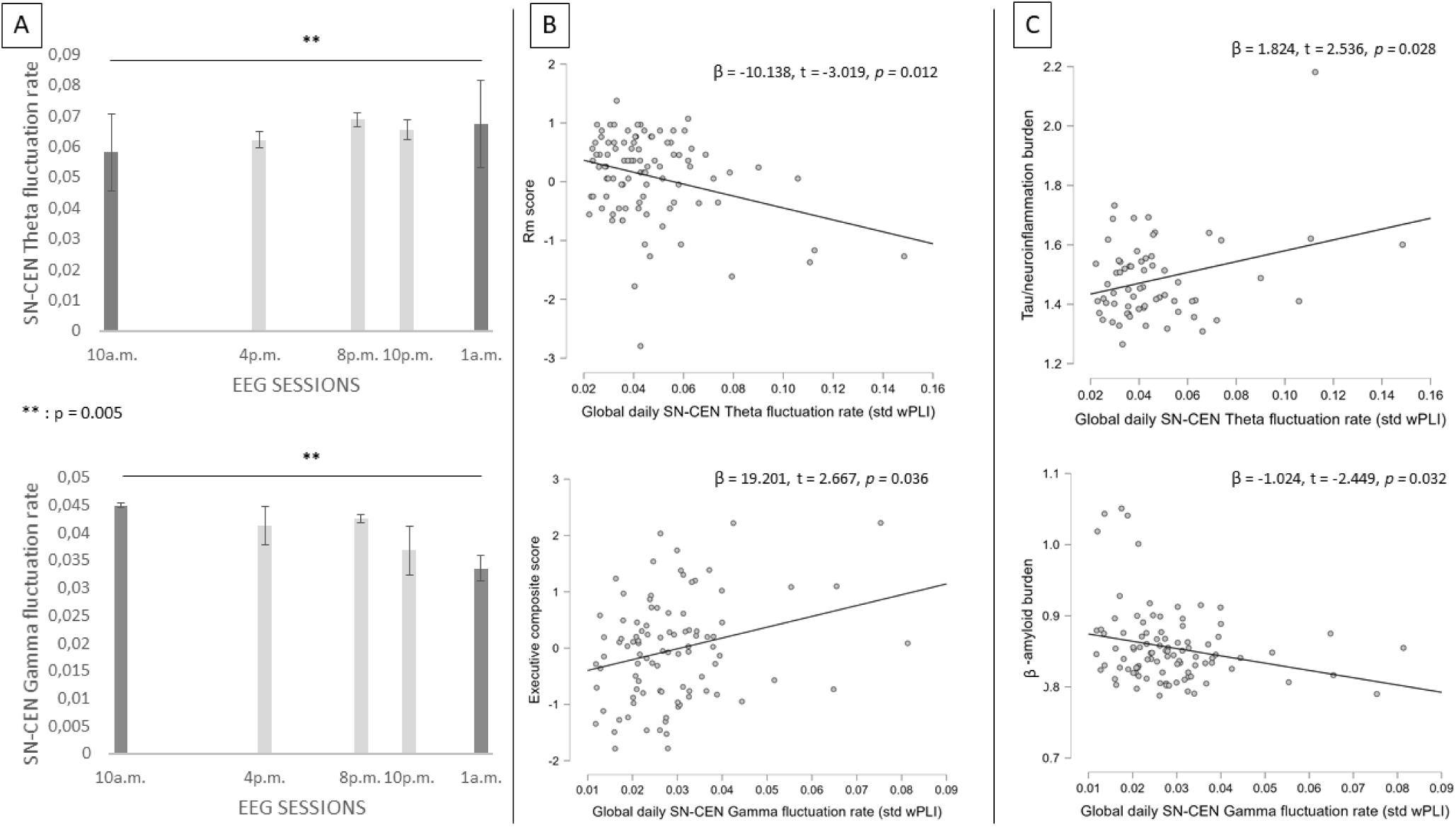
A) Increased Theta and decreased gamma synchrony fluctuations in the SN-CEN inter-k between 10 a.m. and 1 a.m.; B) Negative association between global daily theta fluctuations rate and Rm score (β = –10.138, t = –3.019, *p =* 0.012) and positive association between global daily gamma fluctuations rate and composite executive score (β = 19.201, t = 2.667, *p =* 0.036); C) Positive association between global daily theta fluctuations rate and tau/neuroinflammation burden (β = 1.824, t = 2.536, *p =* 0.028) and negative association between global daily gamma fluctuations rate and β-amyloid burden (β = –1.024, t = –2.449, *p =* 0.032).

### 3.2. Distinct association of theta and gamma SN-CEN inter-network daily fluctuations and cognitive performance

#### 3.2.1. Negative association between Theta SN-CEN daily fluctuations and memory performance

Regression analyses showed that the global theta daily fluctuation rate across the five sessions in SN-CEN inter-network was negatively correlated with the recognition memory score of the Mnemonic Similarity Task (β = –10.138, t = –3.019, Benjamini-Hochberg adjusted p-value = 0.012). These results indicated that higher global fluctuation rates across the day in theta band, within the SN-CEN inter-network, might be related to lower memory performance.

#### 3.2.2. Positive association between Gamma SN-CEN daily fluctuations and executive performance

Within the same SN-CEN inter-network as global theta daily fluctuations, global gamma daily fluctuation rates were positively correlated with the executive composite score (β = 19.201, t = 2.667, Benjamini-Hochberg adjusted p-value = 0.036). Higher global daily fluctuation rate in gamma band within the SN-CEN inter-network, appears to be related to higher executive performance.

### 3.3. Distinct association of theta and gamma SN-CEN inter-network daily fluctuations and early pathological markers

#### 3.3.1. Positive association between Theta SN-CEN daily fluctuations and tau/neuroinflammation burden

We performed regressions analyses to investigate the association between global theta SN-CEN inter-network daily fluctuation rates and tau/neuroinflammation and β-amyloid burden rates. Results showed that global theta SN-CEN daily fluctuation rate was positively correlated with tau/neuroinflammation burden rate (β = 1.824, t = 2.536, Benjamini-Hochberg adjusted p-value = 0.028). These results indicate that higher tau/neuroinflammation burden rate is associated with lower global theta SN-CEN inter-network daily fluctuation rate.

#### 3.3.2. Negative association between Gamma SN-CEN daily fluctuations and early β-amyloid burden

Regression analyses showed a negative regression between gamma SN-CN inter-network daily fluctuation rate and Amyloid-β burden rate (β = –1.024, t = –2.449, Benjamini-Hochberg adjusted p-value = 0.032). Once again, these results suggest that higher β-amyloid burden rate are associated with lower global gamma SN-CEN internetwork daily fluctuation rates.

Post-hoc sub-group analyses, based on participants’ age group (over 60 vs. under 60 years) were performed to further investigate age-related effects and also control for medial split biases (DeCoster et al., 2011). We observed the same correlation patterns between SN-CEN global daily fluctuations in theta and gamma band, their associated cognitive scores and early pathological markers burdens reported below, exclusively within the over 60 age range group (see supplementary for detailed results).

## 4. Discussion

Our main goal was to characterize the daily dynamics of brain activity fluctuations at rest in aging and their associations with cognitive performance and biological markers related to the neuropathology of Alzheimer Disease (AD). For this, we investigated EEG resting-state rhythms fluctuations recorded five times across the day in a wake-extension protocol, completed in late middle-aged healthy participants. Brain fluctuations were considered instead of the mean activity, as recent studies on BOLD-signal highlighted that brain signal fluctuations were predictive of cognitive performance in a way that mean signal cannot capture (Garrett et al., 2011, 2013, 2014, 2020). Moreover, previous M/EEG works showed an association between the increase of resting-state neural fluctuations with aging and cognitive performance (Courtney & Hinault, 2021; Jauny et al., 2022; Hinault et al., 2021, 2023; Uddin et al., 2020; Kumral et al., 2020).

### 4.1. Increased theta and decreased gamma fluctuation rates across the day

We first showed an increase of theta fluctuations and a decrease of gamma fluctuations over the day (i.e. between 10a.m. and 1a.m) at rest. These results can be understood as the reflection of the interplay between sleep homeostasis and the circadian system which results in a non-linear increase of slow theta rhythms synchrony and a decrease of fast rhythms with time awake (Munn et al., 2017). The fluctuation peak qualitatively observed at 8:00p.m. might reflect the maximal strength of the circadian signal opposing sleep need over the so-called wake-maintenance zone. Increased slow-theta fluctuations and decreased fast-gamma fluctuations induced by the circadian processes could reflect a global slowing of brain fluctuations, which might be interpreted as a global decrease of resting-state functional connectivity during the day. This interpretation is in line with previous studies on BOLD-signal diurnal variations (Orban et al., 2020). Taken together, these results seem to highlight that healthy older people exhibit a similar daily temporal organization of brain rhythms fluctuations involved in resting-state functional connectivity as young individuals.

### 4.2. Daily organization of brain rhythms fluctuations involve inter-network couplings

The daily temporal organization of brain rhythms fluctuations was exclusively observed in the functional connectivity of inter-network couplings. Such result is in line with previous works on healthy aging, which showed stronger inter-network functional connectivity and weaker within-network functional connectivity, also termed as functional dedifferentiation, suggesting a less specialized patterns of functional connections (Damoiseaux, 2017; Wig et al., 2017; Setton et al., 2017; Malagurski et al., 2020). Identified couplings were part of the salience – control executive (SN-CEN) and default mode – control executive (DMN-CEN) inter-networks for the theta band. In the gamma band, couplings from the SN-CEN, default mode – salience (DMN-SN), control executive – visual system (CEN-VS) and control executive – dorsal attention (CEN-DAN) inter-networks were observed. The SN, CEN and DMN networks are all three involved in both theta and gamma fluctuations over the day, suggesting that theta and gamma diurnal functional connectivity fluctuations commonly involve interactions between SN, CEN and DMN. These networks include crucial brain regions for whole brain network connectivity and cognitive functioning efficiency (van den Heuvel and Sporns, 2013). Several studies showed an increased functional connectivity between these networks in elders (Ng et al., 2016; Archer et al., 2016).

Interactions between these three resting-state networks were reported in a triple network model aimed at understanding differences in functional connectivity patterns at rest between healthy and cognitively impaired young adults (Seeley et al., 2007; Uddin, 2015), then replicated to investigate these differences among older individuals (Chand et al., 2017). This model proposes the SN to play a role in switching between the DMN associated with internally directed cognitive activities and the CEN involved in externally directed cognitive functions. This inter-network reorganization of the SN with other networks, including the DMN and the CEN has been proposed as the hallmark of aging (LaCorte et al., 2016). It was proposed that the SN drives the DMN and the CEN during resting-state in healthy old participants showing a normal cognitive functioning while the disruption of the control of SN over the DMN and the CEN was associated with cognitive impairment in patients with mild cognitive impairment (MCI) (Chand et al., 2017). Our results showed that daily fluctuations over the SN-CEN inter-network only, in both theta and gamma band were correlated with cognitive performance, namely the recognition memory score (MST) for theta and the executive composite score for gamma daily fluctuations. Thus, our results might provide further details on how this three-part interaction featuring the SN, the CEN and the DMN, fluctuates across the day in aging. Our results importantly reveal that daily fluctuations between SN and CEN networks seems to play a key role in the heterogeneity of cognitive changes associated with healthy aging.

### 4.3. High global daily theta SN-CEN fluctuations are associated with low memory performance

We observed that higher global theta daily fluctuations within the SN-CEN inter-network showed lower memory composite scores. This concerns the recognition memory (rm) sub-score of the Mnemonic Similarity Task (MST), which has been shown to be a sensitive measure to detect subtle general cognitive decline in aging (Pishdadian et al., 2020; Rizzolo et al., 2021). Theta rhythm is widely associated with different memory component, more specifically spatial working memory (Jones et al., 2005), memory encoding (Tambini et al., 2018), and memory integration and storage (Backus et al., 2016). In aging, theta widespread resting-state synchrony has been linked to individuals’ cognitive and memory decline (Klimesh, 1999; Spinelli et al., 2022). Theta synchrony is predictive of the conversion from healthy aging to mild cognitive impairment (MCI) in a group with subjective cognitive decline (Prichep et al., 2006), and of decline from MCI to Alzheimer disease (AD) (Rossini et al., 2006; Caplan et al., 2015; Babiloni et al., 2021 for a review). Thus, considering global daily fluctuations provide further information about theta mechanisms in cognitive aging, such as higher amount of global theta fluctuations during the day seem to be deleterious for memory abilities.

### 4.4. High global daily gamma SN-CEN fluctuations are associated with high executive performance

On the other hand, within the same SN-CEN inter-network, we observed that participants with higher global gamma daily fluctuations also showed higher executive composite scores. In a state of resting-state wakefulness, previous work showed that gamma rhythm reflects neural bottom-up processing and integration of perceptual information between and within different cortical and subcortical brain regions (Jensen et al., 2014). A positive correlation between high global gamma fluctuation rates over the day at rest and executive functioning suggests a more efficient regulation of information processing and maintenance of wakefulness in individuals with preserved gamma dynamics. Our results are also in line with the literature on resting-state synchrony in healthy aging, which showed a higher synchrony of gamma activity in healthy controls relative to patients with mild cognitive impairment (MCI), interpreted in those studies as a compensatory mechanism (Stam Koenig et al., 2005, Prichep et al., 2006; Pusil et al., 2019; Vecchio et al., 2016). These results are also in line with previous studies led on the same COFITAGE cohort, which have shown that less fluctuations of brain responsiveness across the day, measured as cortical excitability, where associated with lower cognitive performance (Gaggioni et al., 2019; Van Egroo et al., 2019). Thus, our results clarify the dynamics of gamma synchronization especially within the SN-CEN inter-network that is associated with preserved executive performance in terms of fluctuations over the day.

### 4.5. Distinct theta and gamma association with AD pathological markers

Our results also provide new information about the association of EEG markers with early tau/neuroinflammation and β-amyloid burden. In fact, the field of clinical neuroscience of ageing is increasingly focused on targeting the earliest signs of neuropathological markers (i.e. β-amyloid and tau/neuroinflammation) in the absence of Alzheimer disease (AD) cognitive symptoms, in order to predict the progression of AD in very early stages (Spinelli et al., 2022). Importantly, distinct association of theta and gamma were observed with tau and with β-amyloid. Indeed, global theta daily fluctuations were positively correlated with tau/neuroinflammation burden in regions corresponding to Braak stages of early regional tau pathology, while global gamma daily fluctuations were negatively associated with β-amyloid burden in earliest aggregation sites for β-amyloid pathology. The positive association between global daily theta fluctuations and tau/neuroinflammation, is consistent with previous works conducted by Gaubert et al. (2019) investigating EEG resting-state activity between groups of patients with positive or negative tau/neurodegeneration. Their results revealed that theta activity in fronto-central regions was higher in tau/neuroinflammation-positive patients. Consistent findings across studies also highlight that both β-amyloid and tau/neuroinflammation were associated with slowing of brain rhythm activity in aging (Tanabe et al., 2020). Here we replicated the association for theta daily fluctuations only with tau/neuroinflammation, which would indicate that tau is associated with different features of theta brain rhythms.

The link between the global daily fluctuations in gamma band and β-amyloid burden could be in line with results in Alzheimer disease’s mice models studies that have reported an association between gamma band activity and β-amyloid, showing a decreased gamma synchrony within the parietal cortex occurring before β-amyloid deposition (Verret et al., 2012) and a reduced β-amyloid burden induced by multi-sensory gamma stimulation (Martorell et al., 2019; Park et al., 2021). Studies in human aging have only found positive associations between local gamma power and β-amyloid burden in AD patients rather than gamma widespread synchrony (see Babiloni et al., 2021 for a review). Studies in human aging are increasingly carried out to clarify the association between gamma-band and β-amyloid burden as gamma sensory stimulation is considered as a potential therapeutic strategy in AD (see Ko et al., 2022 for a review). We here provide arguments to consider gamma fluctuations as a marker of the progression of β-amyloid deposition.

### 4.6. Limits and perspectives

These findings are constrained by methodological limitations. First, our results were obtained from healthy middle-aged to older participants, with a lower mean age that other cohorts of aging individuals, and whose global cognitive performance is in the upper average range. Therefore, the present results could differ in individuals aged 70 and above. Furthermore, the study did not include a group of young adults, so we were not able to assess group differences relative to younger participants to determine the age-specificity of our results. Further studies will aim at investigating the individuals’ cognitive trajectories by assessing the predictive value of daily theta and gamma EEG fluctuations on longer longitudinal protocols, and the evolution of these daily brain fluctuations with advancing age. Notwithstanding, our study is the first to investigate daily dynamics of brain rhythms fluctuations using EEG time-frequency analyses methods in aging. Our results might indicate the involvement of a same functional inter-network, namely the SN–CEN inter-network, in two neural mechanisms (theta and gamma daily fluctuations) which appeared to be associated with distinct cognitive functions. Noteworthy, we assessed functional brain connectivity through the lens of the phase lag index (PLI) which is a bivariate connectivity measure that quantify the activity between couplings of brain regions in a data-driven approach. Forthcoming research using multivariate analyses might be necessary to replicate our findings taking into account the complexity of functional network interactions (Pusil et al., 2019).

## 5. Conclusion

Brain activity fluctuations, long considered as noise, are now considered as a key marker of age-related cognitive variability in several BOLD signal MRI studies (Garrett et al., 2020) contrasting to the small number of EEG studies in this field. Moreover, no study accounted for the daily dynamics of brain fluctuations in the elderly, while it is acknowledged that cognition varies during the day. Here, we showed that a higher level of theta fluctuations at rest is associated with lower memory performance and higher tau/neuroinflammation burden rates, while higher level of gamma fluctuations is associated with a better executive functioning and lower β-amyloid burden, suggesting that these two rhythm fluctuations have specific and opposite roles in cognitive functioning and AD markers progression. Moreover, daily fluctuations in both theta and gamma bands were observed over SN–CEN inter-network, which is in line with the dedifferentiation of network functional connectivity assumed to be a hallmark pattern of aging (Malagurski et al., 2020). Thus, investigating daily EEG fluctuations at rest contributes to better understand subtle brain changes underlying cognitive functioning in healthy aging, in the drive to precise neural fingerprints derived from wake resting-state EEG associated with cognition traits (Castanheira et al., 2021). Results also provide arguments for considering time-of-day when assessing cognition for elders. Using recent EEG frequency analysis techniques enables the identification of early markers of cognitive decline, even with a limited number of electrodes (Gaubert et al., 2020), thereby showing promising potential application for clinical use.

## Acknowledgements

This research did not receive any specific grant from funding agencies in the public, commercial, or not-for profit sectors. Data collection and sharing for this project were provided by the GIGA-CRC in vivo Imaging. Funding for this project was provided by Fonds National de la Recherche Scientifique (FRS-FNRS, FRSM 3.4516.11, EOS Project MEMODYN No. 30446199; Belgium), the Wallonia-Brussels Federation (Grant for Concerted Research Actions—SLEEPDEM), University of Liège, Fondation Simone et Pierre Clerdent, European Regional Development Fund (ERDF, Radiomed Project).

