## Supplementary material for "Daily dynamics of resting-state EEG theta and gamma fluctuations are associated with cognitive performance in healthy aging"

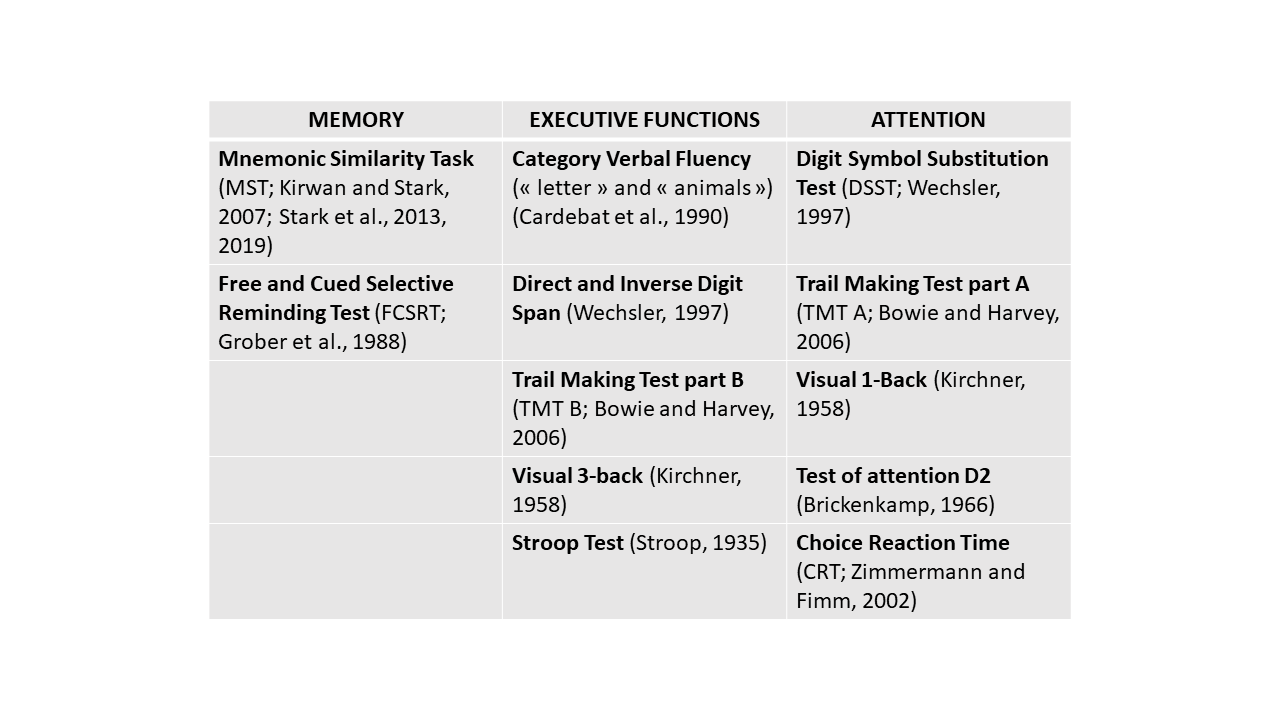
**Table A.** Detailed cognitive assessment of the three specific domains: memory, attention, executive function

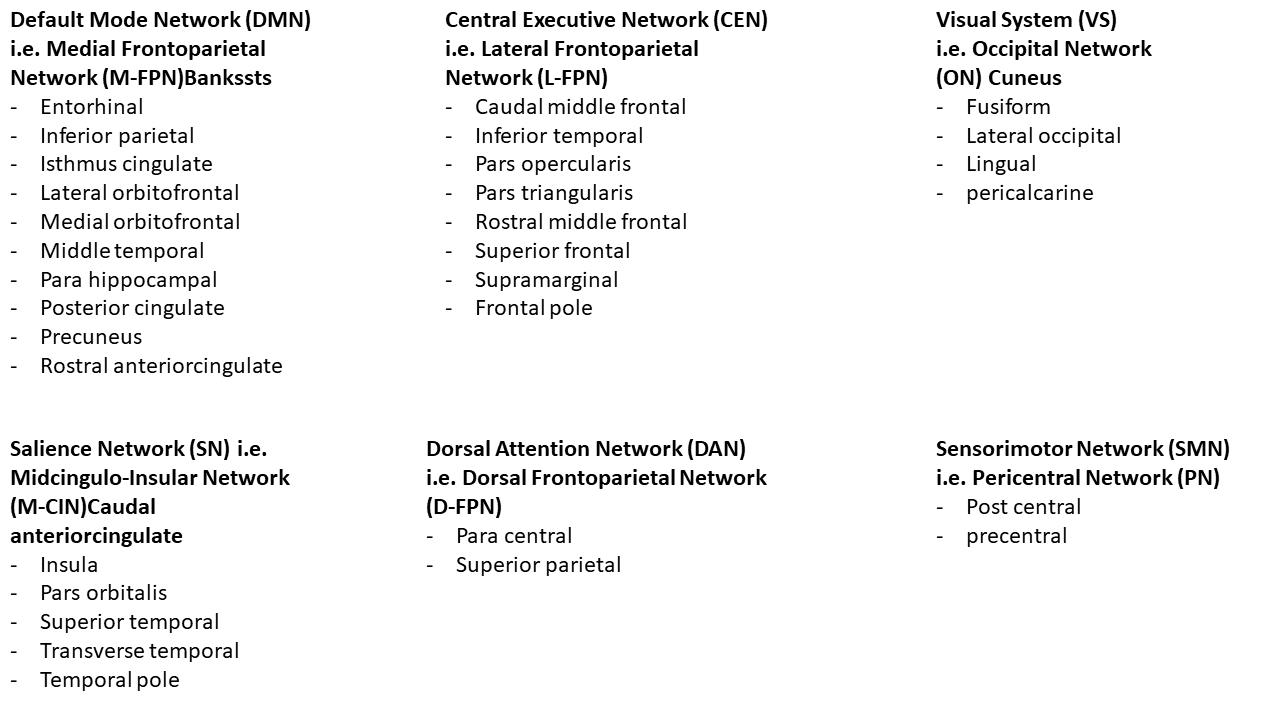
**Table B.** Desikan-Killany ROIs’ attribution to the six Resting State Networks defined by Uddin et al. (2019)

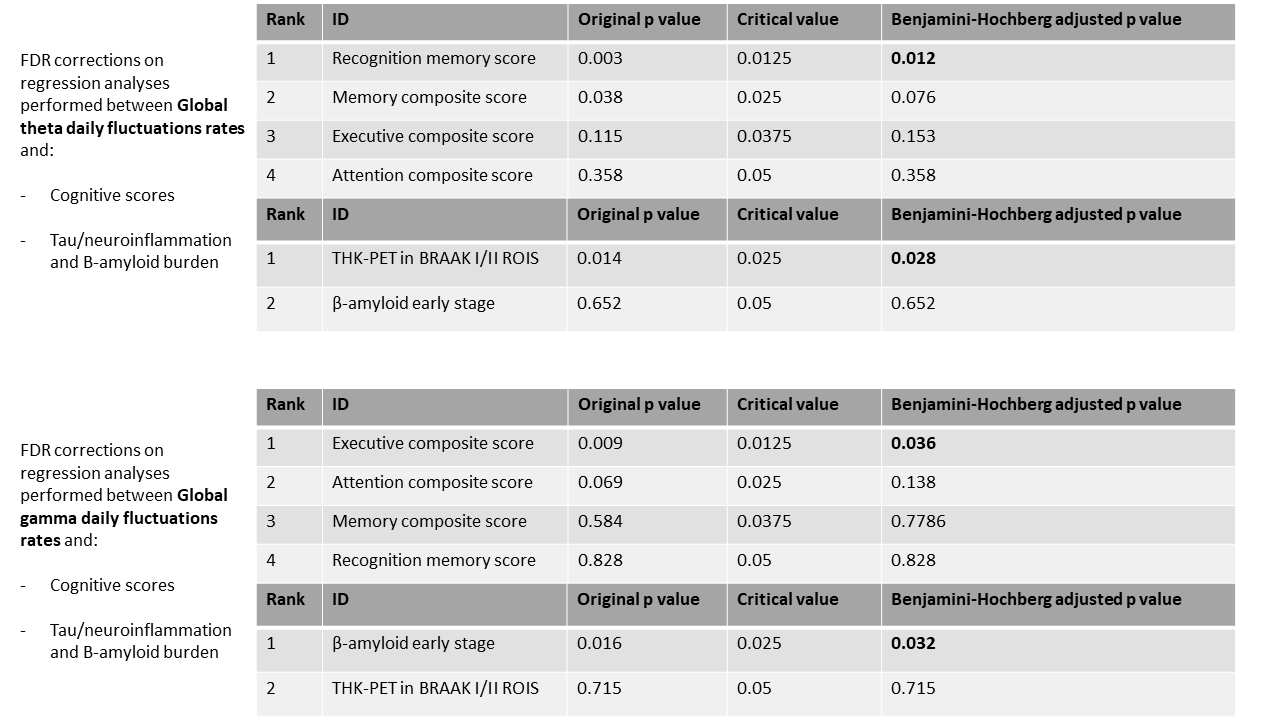
**Table C.** Initial results corrected using FDR (Benjamini and Hochberg, 1995)

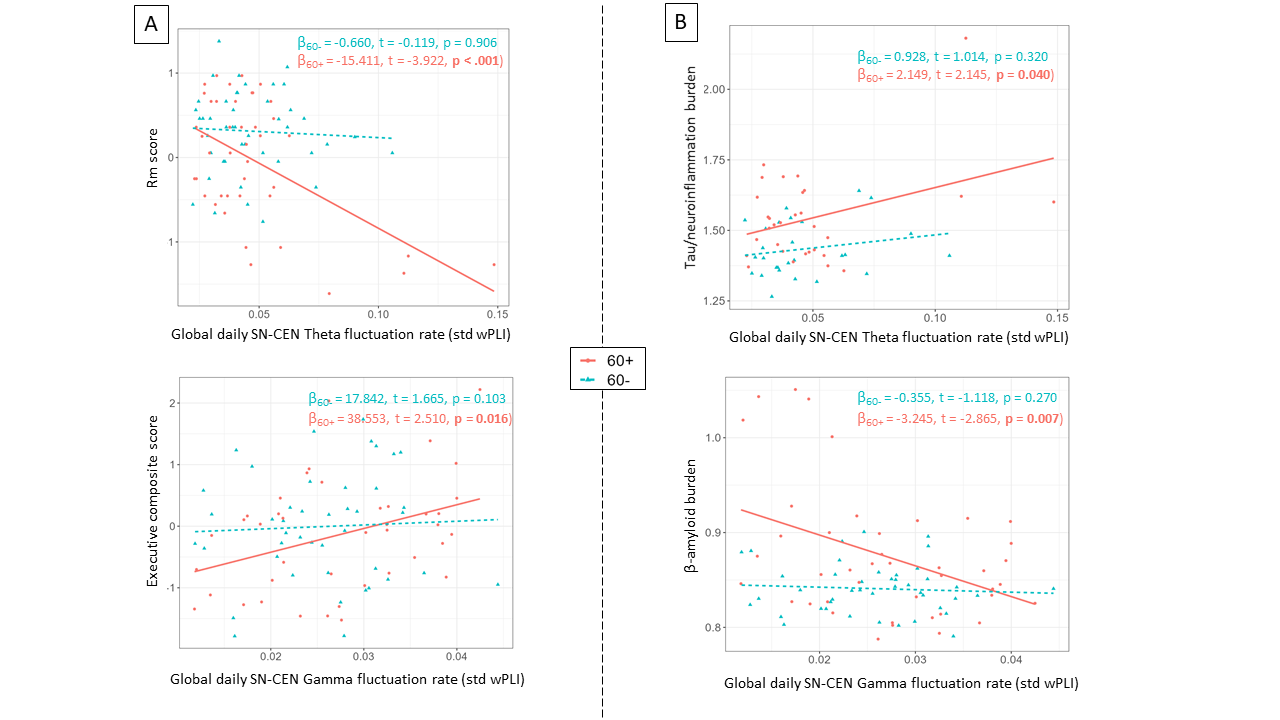

**Figure A.** Post-hoc sub-group analyses based on participants’ age group (over 60 vs. under 60 years). A) Negative association between global daily theta fluctuations rate and Rm score (β = -15.411, t = -3.922, *p < .001*) and positive association between global daily gamma fluctuations rate and composite executive score (β = 38.553, t = 2.510, *p = 0.016*) in over 60 years participants; B) Positive association between global daily theta fluctuations rate and tau/neuroinflammation burden (β = 2.149, t = 2.145, *p = 0.040*) and negative association between global daily gamma fluctuations rate and β-amyloid burden (β_60+_ = -3.245, t = -2.865, *p = 0.007*) in over 60 years participants.
